# Kisspeptin modulates sex-like phenotypes in the neonatal period

**DOI:** 10.1101/2023.11.23.568489

**Authors:** João R. Neves, Miguel Castelo-Branco, Joana Gonçalves

## Abstract

In recent years, our understanding of the physiological role of kisspeptin (Kiss1) has evolved significantly, revealing its pivotal role in shaping sexual differentiation within the brain. Predominantly expressed in the hypothalamus, Kiss1 plays a crucial role in regulating release of the gonadotrophin-releasing hormone (GnRH). Kisspeptin 1 receptor (Kiss1R) exhibits sex-dependent expression in various brain regions, including the medial amygdala and preoptic area (POA) of the hypothalamus, regions underlying sexual development. Importantly, in the critical period for the sexual development in mammals occurs a synchronized surge of Kiss1 and testosterone, highlighting its importance in sexual dimorphism. It has been suggested that the neonatal Kiss1 surge is associated with the buildup of gender-linked social development. We investigated this hypothesis by testing the effect of neonatal blockage of Kiss1 signaling on the sex-like phenotype in rats. We postulate that Kiss1 actions are both time and region-dependent, particularly during the neonatal period, having lasting impact on social interactions.

For that we injected intracerebroventricular (ICV) a Kiss1R antagonist, Kp234, in the first 24h of life. Our data showed, that neonatal Kiss1 is determinant in modulating physiological responses, such as weight and testis development in males (N: males Veh = 10; males Kp234 = 10). Also, we document, for the first time, that Kiss1 modulates the testosterone surge in males and inhibits it in females (N: males Vehicle = 3; males Kp234 = 4; females Vehicle = 3; females Kp234 = 4). Furthermore, neonatal Kiss1 blockade modulated the sexual drive in, while the sexual performance in both sexes is maintained males (N: males Vehicle = 8; males Kp234 = 9; females Vehicle = 7; females Kp234 = 8).

In conclusion, this study underscores the significance of Kiss1 during the neonatal period, a critical phase for sexual differentiation and neuroplasticity. Neonatal Kiss1 emerges as a key player in the development of a sex-like phenotype in rats. Additionally, our findings show dimorphic patterns in Kiss1 influence. This work raises the intriguing possibility that the early influence of Kiss1 in brain circuitry is pivotal in the modulation of gender and plays a role in the social construction of the bimodality of gender identity.

## 1. Introduction

Over the past few years, the knowledge about the biological role in kisspeptin (Kiss1) has advanced namely in shaping the sexual differentiation of the brain (Lee et al., 2022). This peptide family shows predominant expression in the hypothalamus, wherein these peptides play a crucial role in regulating gonadotrophin-releasing hormone (GnRH) (Lee et al., 2022; Oakley et al., 2009). Notably, Kiss1 neurons in the anteroventral periventricular nucleus (AVPV) and arcuate nucleus (ARC) are regulated by estrogen and estradiol, serving as pivotal elements inside the sexual hormone feedback loop (Oakley et al., 2009). Beyond this, the kisspeptin 1 receptor (Kiss1R) is expressed in a sex-dependent manner in other brain areas, such as the medial amygdala and preoptic area (POA) of the hypothalamus (Muñoz de la Torre et al., 2022). Hence, research has demonstrated that Kiss1-Kiss1R signaling is correlated with behavioral dimorphism throughout the lifespan, mainly regarding sexual behavior (Mills et al., 2019).

An especially intriguing discovery is the synchronized rise of Kiss1 and testosterone throughout the neonatal surge, a critical period for sexual differentiation in all mammal species (Jayasena et al., 2014). This temporal contingency underscores the significant function of Kiss1 in sexual dimorphism and differentiation (Mills et al., 2019). Several studies have established that Kiss1R knock-out (KO) displays impaired sexual behavior in adult males, which is restored after testosterone injection. Additionally, Kiss1R is crucial for a proper lordosis response, although it operates independently of GnRH signaling. It is worth noting that plasma testosterone levels are unaltered in neonatal Kiss1R KO adult males, implying that additional factors contribute to the real impact of Kiss1 during the critical neonatal period (Kauffman et al., 2007; Poling et al., 2012). Furthermore, the existing literature predominantly focuses on the effects of Kiss1 on sexual behavior in adult KO mice, which neglects the natural fluctuations of Kiss1 throughout life, such as, the high levels present during the neonatal phase and its near absence in adulthood (Lee et al., 2022).

Importantly, this surge in rodents is proven to be responsible for physiological and behavioral dimorphisms (Lenz and McCarthy, 2010). In humans, this correlation is hard to identify. Most of the sexual differentiation of the brain occurs in the prenatal period, but the effect of postnatal testosterone surge is still understudied (Swaab and Hofman, 1988). Recent studies correlated this surge in humans with gender-linked social development (Alexander, 2014). Additionally, human neuroimaging studies have already demonstrated that Kiss1 administration in adults increases sexual drive, rather than overt physiological responses. Therefore, we hypothesized that Kiss1 can be determinant to develop the neuronal social circuits that will stablish a gender/sex identity and its interactions with the social background, rather than modulating expression of sexual characteristics by itself.

Here, we demonstrate that the actions of Kiss1 are both time and region-dependent, suggesting that neonatal Kiss1 plays a specific role in shaping gendered circuitry development, consequently, having lasting effects on adult social interactions in a sexual behavior context. The study of Kiss1 effects on the onset of the neurodevelopment of gender can imply a scientific paradigm shift. Moreover, these findings have potential societal impact and may contribute to destigmatization of gender. In our research, we intended to give the first steps to understand the nuanced effects that Kiss1 has in the behavior of mammals in the neonatal period.

## 2. Methods

### 2.1 Animals

Females and males Wistar rats were initially purchased at Charles River Laboratories (Germany). Upon arrival animals were kept at standard conditions at the animal facility of Institute of Nuclear Sciences Applied to Health (ICNAS), University of Coimbra. After acclimatization, the breeding process was initiated. Females were allowed to deliver naturally, with the day of birth designated postnatal day 0 (PND0). Immediately, after birth, neonatal animals, within first 24h of life, were submitted to an ICV injection. Four experimental groups were established based on the sex and procedure: males treated with Vehicle (Veh) males treated with Kp234, females treated with Veh and females treated with Kp234. Experimental group formation was blind, and the litters were equally divided between the groups, to reduce bias. All pups were housed together with the mother until PND21. After that, each litter was segregated by sex and treatment. All procedures and experimenters were authorized and complied with the European Union Council Directive (2010/63/EU), National Regulations, DGAV, and ORBEA board of ICNAS (5/2021).

### 2.2 Intracerebroventricular injection

In the first 24 hours after birth, male and female rats underwent ICV injection. One by one, animals were crio-anesthetized. When they became blueish and presented no signs of paw reflex, animals were placed in the stereotaxic apparatus with the head between the ear bars (reversed), parallel to the x-axis, and the bregma-lambda line placed parallel to the y-axis. The syringe was placed above lambda and X and Y coordinates were put on zero. The stereotaxic coordinates of injection were (±0.90;4.2;1.70) mm, to hit the ventricles. 1μL of the solution was slowly injected and the needle was kept in place for around one minute. This procedure is repeated in a bidirectional way. Neonatal animals were put on a warming pad until fully recovered, then returned to their mothers and home-cage and were observed until guaranteed proper mother nursing. Five males and six females injected with Veh, and six males and five females injected with Kp234 solution were sacrificed by decapitation, and their blood collected.

### 2.3 ELISA TESTOSTERONE

Plasma was isolated by centrifugation in EDTA-coated tubes and samples were used on the ELISA kit. The manufacturer protocol was followed (ELISA Kit for Testosterone, Cloud-clone Corp., Houston, USA).

### 2.4 Sexual Behavior test

At PND60, animals underwent a 20-min sexual behavior test was conducted within the dark period, one hour after the beginning of the dark phase in a quiet room where animals were placed at least 2 hours before the test for habituation. These tests were recorded for later analysis using two video cameras (LifeCam HD-3000, Microsoft, Redmond, WA, USA) positioned on the room’s ceiling and in a lateral position. Intact male rats were assessed for copulatory behaviors in response to natural cycling and experienced female rats, whenever they were receptive (previously tested with a 2-minute interaction with an experienced male). The behaviors recorded included the count of mounts and intromission-like behaviors. Additionally, the latency to the first occurrence was documented. Female rats with their natural reproductive cycles intact were observed for proceptive behaviors and lordosis when exposed to experienced male rats. The observations were made daily throughout the estrus cycle, which typically lasted for 5 days, with each observation session lasting 20 minutes. The observed behaviors included proceptive behaviors such as hops (brief jumps with a crouched stance), darts (short sprints followed by a crouched position), and solicitations (facing the male before moving away and adopting a presenting posture). Additionally, lordosis was assessed in response to male rats actively mounting the females. After performing this test animals were sacrificed by decapitation, their weight and length were measured. Also, testicles were extracted and weighed.

### 2.5 Statistical Analysis

The statistical analysis was performed using IBM SPSS Statistics (version 27) at a significance level of p < 0.05. Data distribution was assessed using the Shapiro-Wilk test, while Levene’s test was used to test for homogeneity of variance. Normal distribution was verified, parametric tests were employed, and the results were reported as x̄ = mean ± standard error of the mean (SEM). When comparing two groups, we used a two-tailed independent samples t-test (a Welch’s correction was applied if homoscedasticity was not met). To compare more than 2 groups the results were analyzed with two-way ANOVA followed by Tukey’s multiple comparisons test for sex and treatment differences. Graphs were constructed on GraphPad Prism version 8.0.1 (GraphPad Software, San Diego, CA, USA).

## 3. Results

To evaluate if Kp234 effectively blocks KP signaling, a set of animals (PND0) was sacrificed after 1 hour of injection and plasma levels of testosterone were assessed (Fig. 1A). Regarding males, there was an expected reduction of testosterone levels between male Veh (1.268±0.0349 pg/ml) and male Kp234 (1.073±0.0639 pg/ml; p-value=0.0053). On the contrary, in female groups we found a significant and surprising increase in plasma levels of testosterone (female Veh: 0.6342±0.0301 pg/ml; female Kp234: 0.8018±0.0508 pg/ml; p-value=0.0040). Also, between sexes treatment differences were found. On one hand, the expected increase in testosterone levels in males Veh compared with females Veh (male Veh: 1.268±0.0349 pg/ml; female Veh: 0.6342±0.0301 pg/ml; p-value<0.0001). On the other hand, an unpredicted decrease in males Kp234 compared with females Kp234 (male Kp234 1.073±0.0639 pg/ml; female Kp234: 0.8018±0.0508 pg/ml; p-value<0.0001).

**Fig. 1.**
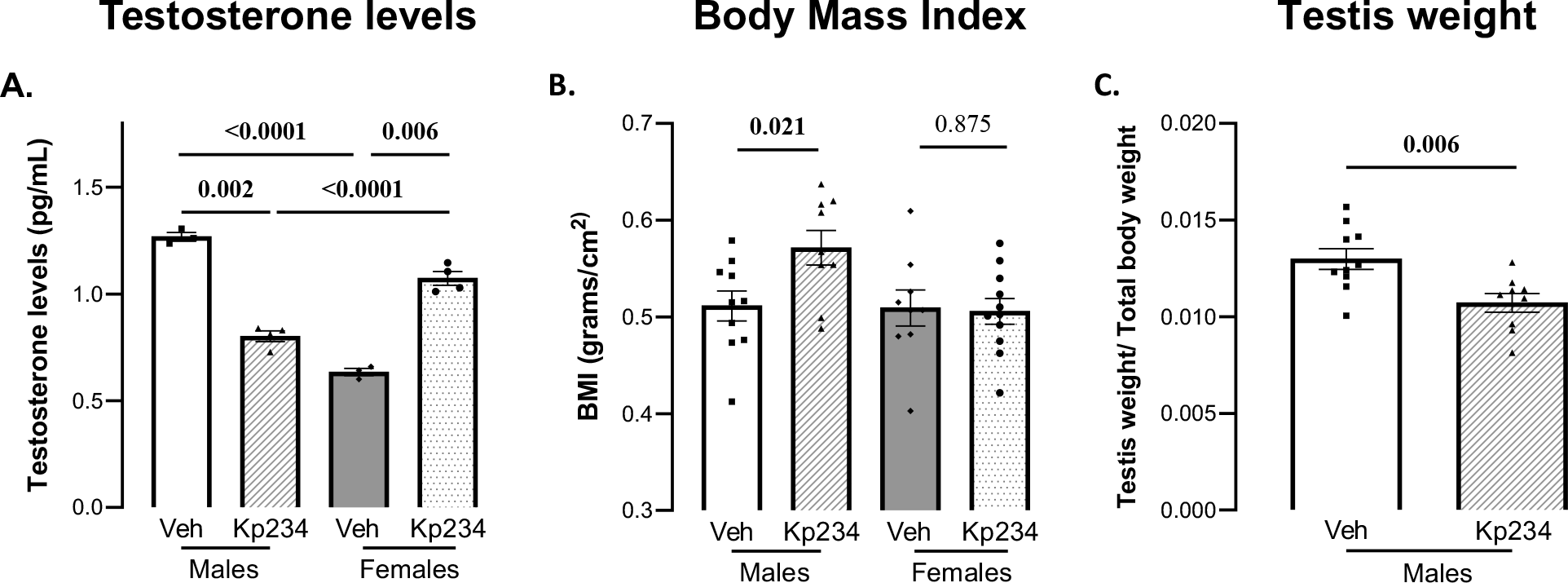
Physiological Measurements. A-Levels of plasma testosterone in sacrificed animals one hour after injection. Two-way analysis of variance; p<0.050; numbers above the bar indicate p-values: in bolt<0.050; males Veh = 3; males Kp234 = 4; females Veh = 3; females Kp234) = 4. B-Body Mass Index (BMI, calculated as the weight division by the square of the length) upon sacrifice. Two-tailed independent samples t-test; numbers above the bar indicate p-values: in bolt<0.050, ns: non-significant (p-value>0.05); males Veh = 10; males Kp234 = 10; females Veh 9; females Kp234 = 11. C-Testis weight normalized by the total weight of male rats. Two-tailed independent samples t-test; number above the bar indicate p-value: in bolt<0.050; males Veh = 10; males Kp234 = 10. Data represented as mean ± SEM.

At adulthood, following behavioral tests, the animals were sacrificed and body mass index (BMI), and testis weight were accessed (Fig. 1B and C). Only males showed significant differences in BMI were males Kp234 presented an increase (0.57±0.018 g/cm^2^) compared with males Veh (0.51±0.015 g/cm^2^; p-value= 0.021). Additionally, we observed that testis weight (corrected to the total body weight) was decreased in males Kp234 (0.011±4.78E-4) compared with males Veh (0.013±5.36E-4; p-value=0.006).

After PND60, when animals reached sexual maturity, sexual behavior was assessed. In male sexual behavior, no differences between experimental groups were observed in both number of mounts and intromission-like behavior (Fig. 2A and C). However, changes were found in latency to start both behaviors (Fig. 2B and D). Indeed, male Kp234 showed a decrease in mount latency (male Veh: 214.9±40.07 sec; males Kp234: 85.18±23.55 sec; p-value=0.013; Figure2B). The same tendency was observed in latency to 1º intromission-like behavior (male Veh: 319.4±51.71 sec; male Kp234 (157.4±37.85 sec; p-value=0.026; Fig. 2D). Concerning female sexual behavior, no statistical differences between experimental groups were detected (Fig. 2E-G).

**Fig. 2.**
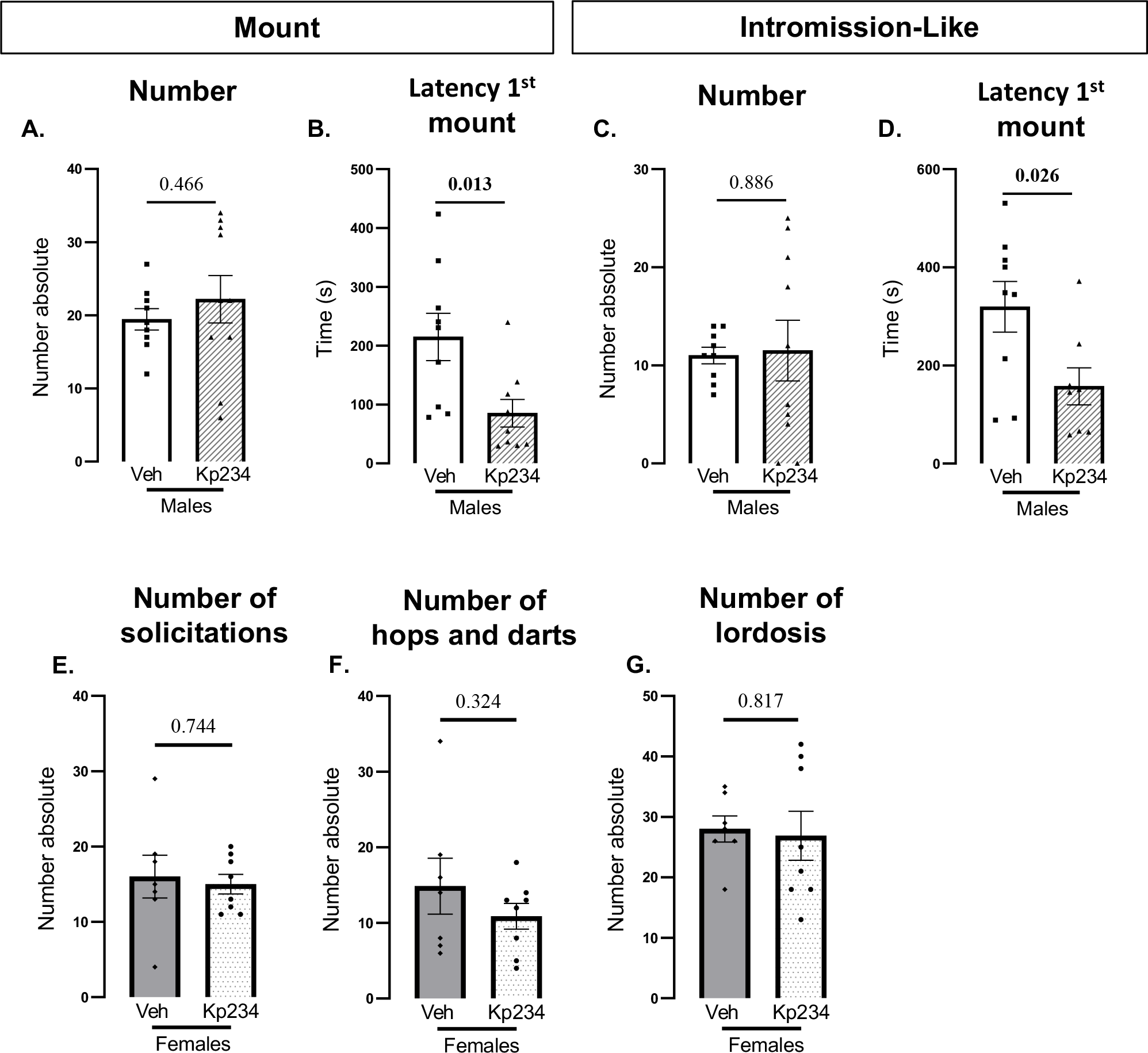
Male and Female sexual behavior after PND60. Male sexual behavior from A until D: A-Number of mounts. B-Latency of the first mount. C-Number of intromission-like behaviors. D-Latency first intromission-like behavior. Two-tailed independent samples t-test: numbers above the bar indicate p-values: in bolt<0.050, ns: non-significant (p-value>0.05); males Veh = 8; males Kp234 = 9. Female sexual behavior from E until F: E-Number of solicitations. F-Number of hops and darts. G-Number of lordosis. Two-tailed independent samples t-test: ns: non-significant (p-value>0.05); females Veh = 7; females Kp234 = 8. Data represented as mean ± SEM.

## 4. Discussion

Here we investigated Kiss1 neonatal signaling as a potential key player in the behavioral shift of the sex-typical behavior. In fact, in rodents neonatal Kiss1 exerts effects on physiologic measures, with a strong sex bias since males showed more dependency of Kiss1 signaling.

Firstly, our results confirmed the notion that Kiss1 modulates metabolism (Harter et al., 2018). This weight variations were already demonstrated to occur due to the direct input of Kiss1 neurons in proopiomelanocortin (POMC) and neuropeptide Y (NPY) neurons in the ARC, with opposite regulation of appetite (Padilla et al., 2010). Importantly, our findings suggested that neonatal Kiss1 blockade could influence appetite or the metabolism across the lifespan in a sex-specific manner. We posited that neonatal Kiss1 antagonist disrupted the usual wiring to POMC and NPY neurons, favoring the increased activity of NPY neurons, which could have led to an increase in feeding. However, more studies are required to better address this hypothesis. Additionally, we observed a decrease in testis weight, which is in accordance with the Kiss1R KO model, underscoring the importance of Kiss1 in gonadal development (Poling et al., 2012). Interestingly, studies have shown that hypogonadism is correlated with increased body weight (Izzi-Engbeaya and Dhillo, 2022).

Despite the extensive literature of Kiss1 effect on metabolism, its importance in modulating the gendered brain remains extensively understudied. A good predictor of Kiss1 influence on gender effects is our finding regarding plasmatic testosterone levels one hour after antagonist injection. We observed that in basal conditions there is the expected increase in testosterone levels between males and females. Further, the expected decrease between males Veh and those injected with the antagonist was also observed, showcasing the influence of Kiss1 in the modulation of the neonatal testosterone surge. However, the important novelty of this work is the finding of an increase in testosterone levels in females compared to males injected with the Kiss1 antagonist and even with Veh. These changes are observed in a period so important for modulating the brain’s sexual differentiation, crucial for the construction of gender behavior. In fact, to our knowledge, there is no literature concerning the influence of Kiss1 on testosterone levels in females neither in the effect of neonatal testosterone on female physiology and behavior. Therefore, more studies focused on studying the behavioral effects of neonatal Kiss1 and its modulation of testosterone levels is crucial to deepen the understanding of how the brain builds the circuitry to develop a gendered phenotype. Interestingly, our data contradicted the evidence found in KO of Kiss1R, where the testosterone surge is maintained after birth (Poling et al., 2012). The differences between our model that transiently blocked the KP signaling with Kiss1R KO model could be due to a compensatory mechanism present in transgenic animals that allowed the recovery of the testosterone peak. Thus, this suggests that this KO model may not be indicated to study the neonatal effect of KP and its influence in building the gendered phenotype.

To explore behavioral influences of neonatal Kiss1 signaling modulation on sex-typical behavior, we examined the sexual performance of both males and females. Importantly, these tests did not evaluate their social response in a context of sexual behavior (Hull and Dominguez, 2007). Therefore, we found no differences regarding female sexual behavior and male number of mounts and intromissions. Probably, its occurred because these behaviors are more related with the sexual performance (Hull and Dominguez, 2007). Nevertheless, in males, the latency to engage in such behaviors presented a significant decrease in males Kp234. These metrics predicted more the willingness to engage in sexual activities and therefore their sexual drive (Hull and Dominguez, 2007). Here, we can conclude that neonatal Kiss1 blockade increased the sexual drive of males. Additionally, a study demonstrated that the masculinization process, mainly the copulatory behavior is more determined by peripubertal Kiss1 modulation rather than the neonatal one (Poling et al., 2012). It seems that neonatal Kiss1 is determinant to control the sexual drive, rather than impact on their sexual performance.

Since sexual behavior is complex and depends on a multitude of different circuitry and regions (Calabrò et al., 2019), we hypothesize that the network responsible for the sexual copulation is Kiss1-independent, at least in the neonatal period. However, sexual motivation seems to be modulated in this time frame. Therefore, it remains unclear what is the role of Kiss1 in the neonatal brain responsible to shape a typical response to external stimuli and how it integrates with the hormonal levels. Therefore, it would be interesting to evaluate the influence of neonatal Kiss1 with different paradigms to probe the social response of both males and females, such as, odor preference paradigms, female sexual motivation in bi-leveled arenas, among others.

In conclusion, this work sheds light, for the first time, on the importance of Kiss1 in modulating the gendered circuitry in the rodent neonatal period. Since there is an overlap with the critical period for sexual differentiation and high neuroplasticity activity. It seems that, in this time frame, Kiss1 is a key player in the buildup of a specific phenotype within the bimodality of gender.

## Acknowledgements

This work was supported by FCT Exploratory project 2022.01066.PTDC and FCT Strategic plan UIDP/04950/2021 (CIBIT).

## Ethical approval statement

All procedures and experimenters were authorized and complied with the European Union Council Directive (2010/63/EU), National Regulations, DGAV, and ORBEA board of ICNAS (5/2021).

## CRediT authorship contribution statement

J.R.N. - Data curation, Formal analysis, Methodology, Software, Visualization, Writing – original draft. M.C.B. and J.G - Conceptualization, Funding acquisition, Methodology, Resources, Supervision, Validation, Writing – review & editing.

